# Probabilistically Weighted Multilayer Networks disclose the link between default mode network instability and psychosis-like experiences in healthy adults

**DOI:** 10.1101/2021.05.17.444398

**Authors:** Simone Di Plinio, Sjoerd J H Ebisch

**Affiliations:** Department of Neuroscience, Imaging, and Clinical Sciences, G D’Annunzio University of Chieti-Pescara, Chieti, Italy; Institute for Advanced Biomedical Technologies (ITAB), G D’Annunzio University of Chieti-Pescara, Chieti, Italy

**Keywords:** Multilayer networks, Dynamic connectivity, Default mode network, Psychotic experiences, Functional networks, Flexibility

## Abstract

The brain is a complex system in which the functional interactions among its subunits vary over time. The trajectories of this dynamic variation contribute to inter-individual behavioral differences and psychopathologic phenotypes. Despite many methodological advancements, the study of dynamic brain networks still relies on biased assumptions in the temporal domain. The current paper has two goals. First, we present a novel method to study multilayer networks: by modelling intra-nodal connections in a probabilistic, biologically driven way, we introduce a temporal resolution of the multilayer network based on signal similarity across time series. This new method is tested on synthetic networks by varying the number of modules and the sources of noise in the simulation. Secondly, we implement these probabilistically weighted (PW) multilayer networks to study the association between network dynamics and subclinical, psychosis-relevant personality traits in healthy adults. We show that the PW method for multilayer networks outperforms the standard procedure in modular detection and is less affected by increasing noise levels. Additionally, the PW method highlighted associations between the temporal instability of default mode network connections and psychosis-like experiences in healthy adults. PW multilayer networks allow an unbiased study of dynamic brain functioning and its behavioral correlates.

## INTRODUCTION

Multilayer networks rely on the assumption that different network layers can be combined in supramodal entities since elements (nodes) occupying the same place in each network can be considered as naturally linked across time or experimental conditions of a task (Hutchinson et al., 2013; Boccaletti et al., 2014; Kivela et al., 2014; De Domenico et al., 2017; Thompson et al., 2018). Defining a supramodal structure allows us to investigate dynamic changes in a network indexed by dynamic nodal measures such as promiscuity, flexibility, integration, and recruitment (Bassett et al., 2011; Papadopoulos et al., 2016; Telesford et al., 2017; Pedersen et al., 2018). Multilayer networks have increasingly been employed in both basic and clinical neuroscience and highlighted interesting patterns of dynamic network rearrangement underlying normal behavior (Braun et al., 2015; Telesford et al., 2016) as well as clinical conditions such as depression (Zheng et al., 2018) and schizophrenia (Gifford et al., 2020).

However, actual models of multilayer brain networks lack an objective methodological procedure to weight network parameters in a biologically-sound fashion. The topological and modular properties of multilayer brain networks rely on two key parameters, namely *gamma* (γ) and *omega* (ω). The former, γ, indicates the weight of the null models when computing modularity (Lancichinetti and Fortunato, 2009, 2012; Betzel and Bassett, 2017). Thus, increasing the value of γ results in a more fragmented network structure with many small modules (Bassett et al., 2013; Gu et al., 2015; Nicolini and Bifone, 2016; Betzel et al., 2017). The latter, ω, represents the empirical strength of connections that a node has toward itself in consecutive (if multilayer networks are built across time) or alternative (if multilayer networks are built across experimental conditions) time intervals (De Domenico et al., 2013; Muldoon et al., 2016). Thus, ω essentially controls the intra-layer versus cross-layer modular architecture. The current standard is to set ω to fixed numbers and to investigate the effect of varying these “fixed-ω” on the modular architecture. However, as already noted by many scientists, such an assumption may introduce strong biases in the analysis since it implies arbitrariness regarding the values of the detection of the modular architecture. For example, by setting ω=1, dynamic (temporal) oscillations of nodal activity over time may be non-efficiently integrated into the cross-layer modular detection (Chai et al., 2016; Betzel and Bassett, 2017).

Many studies that investigated dynamic evolutions of brain activity and functional connectivity highlighted classes of brain regions with peculiar functional properties in the time domain (Van de Ville et al., 2010; De Pasquale et al., 2016; Breakspear, 2017). For example, regions in the inferior parietal lobule, medial premotor cortex, and posterior cingulate cortex have been labelled as functional hubs (De Pasquale et al., 2018) since they dynamically link regions in their modules (cingulo-opercular network, sensorimotor network, and default mode network, respectively) with other brain modules (O’Neil et al., 2015; Preti et al., 2016). The efficient integration of information across networks is crucial for controlling explicit, externally directed actions (Spadone et al., 2019; Wens et al., 2019) and for implicit, self-related information processing (Di Plinio et al., 2020a). Aberrant connectional profiles, especially in regions related to the default mode network, seem to underlie many psychopathological conditions (Brovd et al., 2009; Hua et al., 2019), and these characteristics may depend on dysfunctional dynamic processes (Braun et al., 2015, 2016; Du et al., 2018). Thus, implementing a degree of control over the temporal domain is necessary to increase the ability to detect true transient module configurations during conscious experiences, which are naturally occurring both during task execution and during task-free states (Liu et al., 2020).

Here, we introduce an original parametrization for detecting dynamic modular architectures intending to improve the biological appropriateness of community detection in multilayer networks. As described above, the current estimation of the modularity relies on a structural (γ) and a pseudo-structural (ω) parameter. Instead, the parametrization proposed here accounts for uncertainty in the temporal domain and replaces ω. We introduce beta (β), a parameter that regulates the probability distributions of the strength of edges connecting nodes across multiple layers of a multilayer network (intra-nodal, cross-layer connections). We also implement a biologically driven way to assign strengths of intra-nodal (temporal) connections from these probability distributions by employing spectral coherence (Bowyer, 2016; Mohanty et al., 2020) to choose the appropriate configuration of probabilistic intra-nodal weights. In other words, our method probabilistically regulates intra-nodal weights across layers based on the biological and temporal properties of the node’s signal.

Our approach allows different strengths of cross-layer connections to improve the appropriateness of multilayer modular detection. The procedure mirrors the current methodology for selecting structural resolutions through the parameter γ and aims to investigate modular architectures of the brain by using multiple temporal resolutions and multiple structural resolutions, to implement an unbiased methodology. The probabilistically weighted (PW) multilayer community detection proposed in this paper is expected to increase the accuracy toward the detection of physiologically plausible, dynamic modular structures by eliminating the biases constrained by a fixed and “guessed” value of ω. The modulation of temporal resolution in PW networks allows for uncertainties over temporal multilayer network features in the same way as γ enables the control over spatial network features.

We test this new approach both on simulated data and on real resting-state fMRI acquisitions of 39 healthy participants. The new method’s performance was tested together with standard (fixed-ω) methods. To compare the performances of the methods, we investigated synthetic, weighted networks obtained by varying the number of modules, the weight of structured and unstructured noise in the time series, the structural resolution parameter (γ), and the values regulating the standard parameter (ω) and our proposed parameter for probabilistic weights based on spectral coherence (β). Moreover, the association between multilayer network properties and individual differences in behavior was tested, focusing on the neural correlates of psychosis-relevant personality traits, which have been hypothesized to be related to self-dysfunction and network segregation (Ebisch and Aleman, 2016; Humpston and Broome, 2020; Di Plinio et al., 2020a). Subclinical, psychosis-relevant personality traits were evaluated, which can be detected on a continuum in the general population below any threshold of a clinical diagnosis (McGrath et al., 2015). Such traits can be considered critical from a clinical point of view (Orr et al., 2014), as well as from a phenomenological perspective to provide insight into the relationship between neurobiology and the sense of self (Humpston, 2014).

## METHODS

### Synthetic Networks

Three modules in a network of 100 nodes were simulated as follows in 10 consecutive time windows. Each node was initially assigned to a specific module across all the time windows (30% of the nodes were assigned to module A, 50% to module B, 20% to module C). Then, 40% of the nodes were chosen at random to be fast/slow/random oscillators (15%, 15%, 10% respectively) to simulate diverse physiological properties of brain hubs. Random, slow, and fast oscillators switched between their module and another random module every 1, 3, and 5 layers, respectively.

Once defined the modular time-varying structure, time series were created as follows. First, for each time window, a prototype matrix of synthetic functional connections was simulated as a 100×100 symmetric Lehmer matrix. Since in the Lehmer matrix the values of the functional connections increase with proximity to the diagonal and with progressive node numbers, such matrix allows the generation of modules with different degrees of intrinsic connections, thus, avoiding biases related to connection strengths. A time series was simulated for each node from the starting correlation structure, and, for each module, the nodes’ time series were selectively time-shifted to create modular structures.

Within each node, the time series of 100 voxels were simulated to simulate realistic voxels within a parcel. Initially, for each node, the node’s time series was assigned to each voxel within the node. Then, two sources of noise were simulated to represent physiologically unrelated noise (unstructured noise) and the “interference” of nearby biological units in the node signal (structured noise). Unstructured noise was simulated - for each voxel - as a random time series that was added to the voxel signal in a weighted manner. Structured noise was simulated by choosing a random [2 to 5] number of attractors on the external surface of the node. A random time series was simulated for each attractor. Then, each voxel in the node suffered each attractor’s interference depending on its closeness to the attractor. Finally, to mirror common procedures in neuroimaging, each node time series was obtained by averaging the time series of the voxels within it. Different weights were considered for the two noises (low, medium, high) to investigate the impact of the signal-to-noise ratio to the module detection across the two methods.

Once noisy time series were synthesized, the generalized Louvain function was used to detect multilayer modules following the multilayer network quality function (Mucha et al., 2010) implemented with codes from Jeub and colleagues (“A generalized Louvain method for community detection implemented in MATLAB” http://netwiki.amath.unc.edu/GenLouvain/GenLouvain). The generalized Louvain function was implemented both in the “standard” form (fixed-ω) as well as using the probabilistically weighted method (PW). The whole procedure was repeated for 100 independent cycles so that, in each cycle, a new modular structure was randomly created. The three levels of structured/unstructured noises were introduced for each cycle. As suggested by many relevant studies, it is not only important to test inter-layer connectional strengths, but it is also crucial to vary the structural resolution parameter of brain networks (Chai et al., 2016; Betzel et al., 2019; Puxeddu et al., 2019; Yang et al., 2021; Tardiff et al., 2021). Accordingly, we investigated a range of structural resolutions, modular structures, and inter-layer functional weights in the estimation of multilayer networks. More specifically, the structural resolution parameter (γ) was varied between 0.5 and 2.0 with steps of 0.1. Different values for the parameter establishing fixed cross-layer connections (fixed-ω) were considered ([0.10 0.25 0.50 0.75 1.00]). Moreover, the PW method was also implemented without assumptions about the average or median connectional strengths (see below). Analyses were performed using MatLab version 2019a (The MathWorks). The weighted approach is described in the next paragraphs. The procedure for obtaining synthetic networks is schematised in Figure 1. To assess the independence of the methods’ performances from the number of modules in the network, the whole procedure was repeated using five modules of equal size (20% of the nodes assigned to each module).

**Figure 1.**
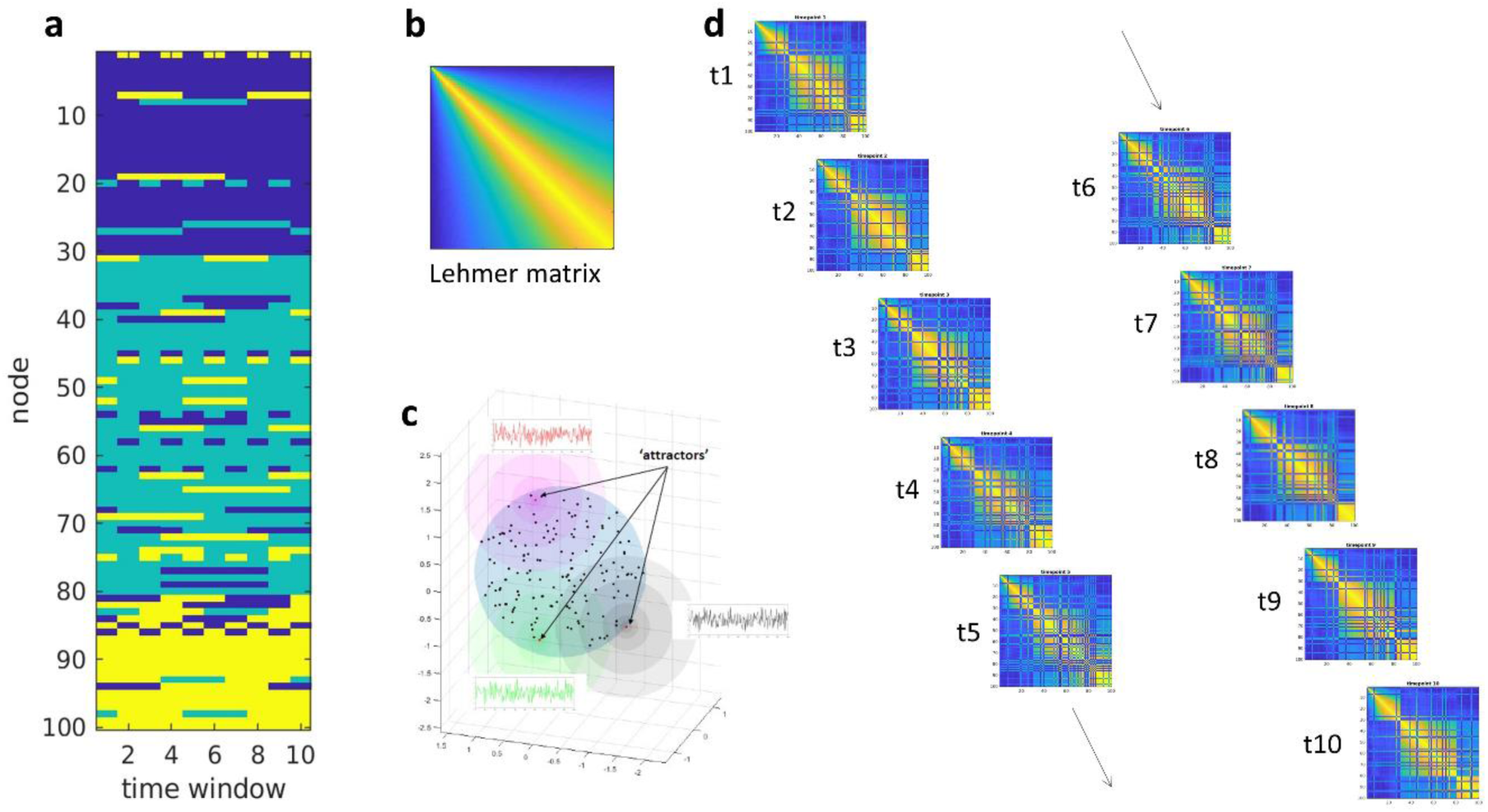
Synthetic multilayer networks. (a) The matrix of synthetic node communities was generated for ten consecutive timepoints. To note, the communities were different at each cycle. (b) A Lehmer matrix was created. Values inside the matrix range from 0 to 1 to simulate functional connectivity among brain regions. A time series was generated for each of the voxels within each of the 100 nodes. Each voxel was affected by unstructured and structured noise. (c) While the unstructured noise was a random time series generated independently for each voxel, the structured noise was specific for each node. The subfigure depicts an example of “noisy attractors” which differentially influenced node’s voxels depending on the distance. (d) For each module, nodes’ time series were selectively time-shifted to create modular structures. After the procedure, at each cycle, a series of 100 timeseries arranged with the desired modular structure and with controlled signal-to-noise ratio was produced. These time series were used as input for the calculation of functional connectivity matrices. The figure illustrates the three-modules structure. The same procedure was used for the five-modules structure.

### Probabilistically Weighted (PW) Multilayer networks

To implement control over nodal temporal features of the network, we introduced a matrix of probability weights W. Values in W were controlled by the temporal resolution parameter β and by the temporal features of the nodes. As probability weights, values in W basically represent the probability of nodes to be self-connected among different layers of the network. Thus, with temporal multilayer networks, W is a N by T-1 matrix in which T is the number of time windows and N is the number of nodes in the network. By contrast, if working with task multilayer networks, the matrix W would be a N by K matrix in which K represents within-node cross-condition interconnections and is calculated as K = C! In which C is the number of experimental conditions (e.g., in a classic two by two factorial psychology paradigm, there would be four conditions, so that K = 4! = 24).

To control the strength of cross-layer connections and to mirror the weight of the null model for the structural resolution parameter, values of W were defined using a probabilistic approach. Initially, a set of Levy alpha-stable distributions, Ω, was generated. Stable distributions rely on four parameters, allowing complete control over the shape and the scale of the distribution. The rationale behind using stable distributions is to have reasonable values of (positive) cross-layer functional connections between 0.0 and 1.0 which symmetrically vary with a center in 0.5, reflecting the situation in intra-layer functional connections usually derived using Pearson correlations. The beta parameter is particularly convenient since it allows to regulate left-skewness and right-skewness of the distributions symmetrically. To reflect correlation (functional connectivity), the values of W (probabilities) were sampled within the interval [0 1]. Thus, the location parameter of the distributions, μ, was set to 0.50 and the scale parameter, σ, was set to 0.75. The first shape parameter, α, was empirically set to 0.40, while the second shape parameter, β, was modulated within the range [-1 1] to obtain a series of distributions in which the head and the tails are arranged progressively in the interval [0 1]. In other words, a set of probability distributions (Ω) was created in which the ratio of low vs high probabilities was controlled through the parameter β: with higher, positive values of β, the head and the fat tail of a specific distribution in Ω are located toward lower probability values (lower probabilities of cross-layer connections); instead, with lower, negative values of β, the head and the fat tail of a specific distribution in Ω are located toward higher probability values (higher probabilities of cross-layer connections). Values used for β in this study were: [-.75 -.25 0 .25 .75].

The distributions in Ω were used to fill values in W based on the coherence between time intervals. More specifically, we used spectral coherence, also known as magnitude-squared coherence (Bowyer, 2016; Mohanty et al., 2020), across consecutive time windows of each node. Coherence estimates the consistency of amplitude and phase between signals across frequencies, and thus it is biologically appropriate grasping coherent signal fluctuations of a node in two adjacent time windows (or in two experimental conditions) *i* and *j*. In other words, coherence is appropriate to assess if two temporal fragments related to the same node have low or high probability to be connected – i.e., to be “self-similar” – between the layers *i* and *j*. We would like to note that many other alternatives to the coherence, like the simple correlation, would be unsuitable except in the case in which temporal segments are aligned, or time-locked to specific stimuli. This only occurs with task experiments using block-designs, with blocks of even length. Instead, coherence works in the frequency domain and not in the time domain, and thus is valid in any experimental design. To have an approach which does not theoretically depend on the way data was acquired is a major advantage and is guaranteed only by using the coherence. Using this procedure, the temporal resolution, β (the β shape parameter of the stable distributions), regulated the probabilities of connections among adjacent time windows in a way driven by the similarity of the node’s signals in the two consecutive time intervals. The procedure for estimating probabilistically weighted multilayer modules is illustrated in Figure 2.

**Figure 2.**
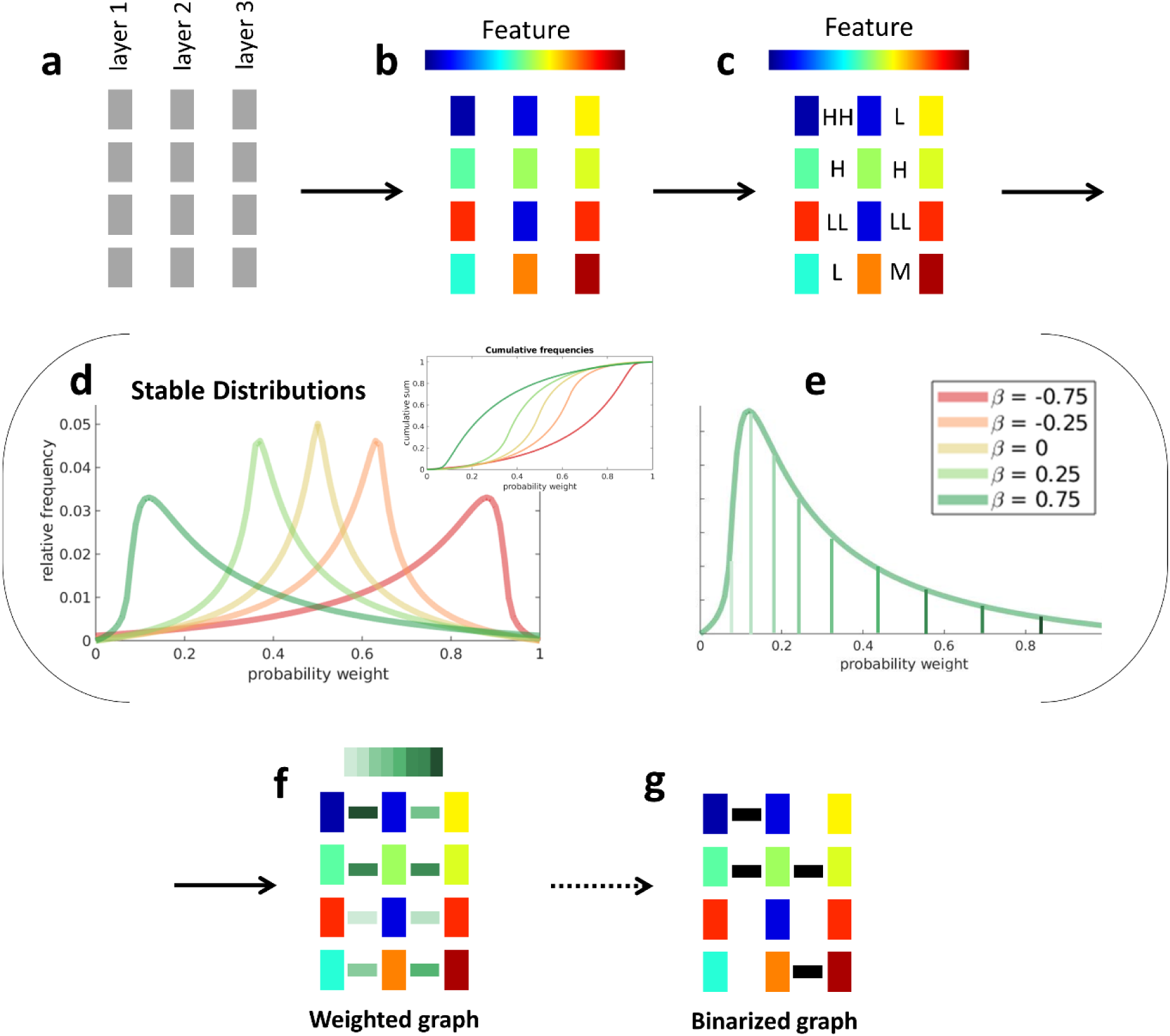
Toy example with 4 nodes and 3 layers (a), illustrating the procedure for estimating probabilistically weighted cross-layer links. Temporal (biological) features (e.g., frequencies amplitudes) are extracted for each node, in each layer (b). The similarity of temporal features is calculated for each adjacent layer (c) using coherence, producing a N x L-1 matrix, where N is the number of nodes and L is the number of layers (HH=very high similarity; LL= very low similarity). Concurrently, a vector of N x L-1 probability weights, W, is obtained from a sample distribution for each value of β considered in the probabilistic analysis (d). For example, with β = 0.75, most links will have a value between 0.1 and 0.2 (e), and high similarities will be associated with higher values. Then, weights are assigned according to both the values in W and the signal similarity over time (coherence), so that cross-layer links connecting nodes with high self-similarity over time are higher (f). Weights can be eventually used as probabilities for the linkage to occur in binary networks (g).

### Statistical Analyses of Simulated Data

The performance of the two methods was assessed by calculating the degree of similarity between the true simulated modular structure and the estimated modular structure through two commonly used indexes for calculating correlations across modular structures, namely the adjusted mutual information index (AMI) and the Rand coefficient. These indices are commonly employed in research and indicate the performance of a modular detection (i.e., clustering) procedure when the ground truth is known (Vinh et al., 2010).

### Application on resting-state data

The procedure described above for detecting PW multilayer networks, together with the standard ω=1 procedure was implemented also on resting-state data from a sample of 39 healthy participants (19 females and 20 males, aged 23 ± 2; 35 right-handed and 4 left-handed) without a history of psychiatric or neurological disease and contraindications for MRI scanning participated in the experiment. The experiment was approved by the local ethics committee (Comitato Etico per la Ricerca Biomedica delle province di Chieti e Pescara) and by the Department of Neuroscience, Imaging and Clinical Sciences (DNISC) of the G. d’Annunzio University of Chieti-Pescara. All participants had a normal or corrected-to-normal vision and provided written informed consent before taking part in the study in accordance with the Declaration of Helsinki (2013).

### Resting-state Data acquisition

Each participant performed two consecutive task-free fMRI runs, each consisting of 376 volumes. The participants were instructed to watch a white fixation cross on a black screen without performing a cognitive task. Each run lasted approximately 7.5 minutes. Functional images were acquired using a Philips Achieva 3T scanner installed at the Institute for Advanced Biomedical Technologies (Gabriele d’Annunzio University, Chieti-Pescara, Italy). Whole-brain functional images were acquired with a gradient echo-planar sequence using the following parameters: repetition time (TR) = 1.2 s, echo time (TE) = 30 ms, field of view = 240×240×142.5 mm, flip angle = 65°, in-plane voxel size = 2.5 mm^2^, slice thickness = 2.5 mm. A high-resolution T1-weighted whole-brain image was also acquired after functional sessions using the following parameters: TR = 8 ms, TE = 3.7, FoV = 256×256×180 mm, flip angle = 8°, in-plane voxel size = 1 mm^2^, slice thickness = 1 mm.

### Resting-state Data analysis

Functional connectivity was calculated as the correlation among average parcels timeseries for a total of 418 nodes using cortical and subcortical atlases from Joliot and colleagues (Joliot et al., 2015) plus the cerebellar atlas from Diedrichsen and colleagues (Diedrichsen et al., 2009). These atlases were chosen because of the integrative method used to define both cortical and subcortical parcels without lateralization biases. The functional connectivity values, that is, the edges of the connectomes, were obtained using the z Fisher transform of the Pearson correlation among pre-processed time series and were used to create individual (subject-specific) weighted graphs. The modular architecture was visualized using BrainNet Viewer (www.nitrc.org/projects/bnv/). We used both the PW method and the standard “fixed-ω” implemented to detect time-varying modular structures across 10 consecutive, non-overlapping time windows of equal length. For consistency, the procedure implemented for the analysis of multilayer networks in real data was the same as described above for the simulations.

We tested if brain-behavioral resting-state associations were differentially detected by the standard approach with fixed-ω and by the PW method. Thus, nodal metrics of flexibility, integration, promiscuity, and recruitment (Bassett et al., 2011; Papadopoulos et al., 2016; Telesford et al., 2017; Pedersen et al., 2018; Sizemore and Bassett, 2018) were estimated for each subject using both the fixed-ω and the PW methods. The *flexibility* of each node corresponds to the number of times that it changes module allegiance, normalized by the total possible number of changes. The *integration* coefficient of a node is the average probability over time that it is in the same community as regions from other systems. The *promiscuity* of a node indicates the fraction of all the communities in the network in which the node participates at least once. Finally, the *recruitment* coefficient corresponds to the average probability that a node is in the same network community as other regions from its own system.

The behavioral variables were represented by five latent, orthogonal psychosis-relevant factors that have also been used in previous works to study the sense of self and the sense of agency (Di Plinio et al., 2019, 2020a, 2020b). We investigated the relation between dynamic multilayer metrics and psychosis-related personality traits with a focus on factors related to psychotic disorders (Asai et al., 2011; Gallagher et al., 2016). These five factors were reliably (Tucker congruence coefficient = 0.95) derived from a factor analysis with orthogonal (orthomax) rotation on a series questionnaires that were administered to a larger sample (N=101) including the present fMRI sample (see Di Plinio et al., 2019, 2020a, 2020b for the complete factor analysis procedure) The questionnaires measured basic as well as psychosis-relevant personality traits: the Big-Five Questionnaire (BFQ short version, Soto and John, 2017), the tolerance of uncertainty scale (IUS-12, Carleton et al., 2007), the Schizotypal Personality Questionnaire (SPQ, Raine et al., 1991), the Community Assessment of Psychic Experience (CAPE, Konings et al., 2006), and the State-Trait Anxiety Inventory (STAI2, Spielberger et al., 1983). The five factors showed the highest loadings in the following subscales: factor 1: STAI2, IUS-inhibitory, CAPE-depression, BFI-neuroticism (negative affect); factor 2: SPQ-cognitive perceptual, CAPE-positive, CAPE-negative (psychosis-like experiences); factor 3: IUS-prospective, BFI-extraversion, BFI-agreeableness, BFI-openness (sociality); factor 4: SPQ-interpersonal, SPQ-disorganizational (schizotypal); factor 5: BFI-conscientiousness.

### Statistical Analyses of Real Data

Brain-behavioral relationships were investigated using whole brain, univariate mixed-effects models in which the five psychometric factor scores were used as continuous predictors for each multilayer metric of interest; random intercepts and slopes were added at the individual node level to allow for ROI-specific weighting of brain-behavior correlations; random intercepts were added at the subject level. To properly assess meaningful brain-behavior correlations, we implemented a cross-γ comprehensive regression model by using γ as a further random grouping factor. For this analysis, γ was varied in the interval [0.6 1.5] with steps of 0.1. A separate model was built for each method (fixed-ω, PW) and for each parameter value used for detecting multilayer networks and related theoretical measures. Major nodal contributions in the brain-behavior associations were detected using best linear unbiased predictors (BLUPs) to generate nodal conditional expectation (ICE) plots and to highlight nodes with the highest contribution.

## RESULTS

### Simulations

The PW method outperformed the fixed-ω method by far as demonstrated by differences in the indices used to measure the correspondence between true modular structures and reconstructed modules (i.e., adjusted mutual information and Rand coefficient). The PW method outperformed the standard method with every noise combination and independently of the structural resolution parameter γ.

Regarding the three-modules structure, as shown in Figure 3a, the effect of the term *method* in the models was significant across structured and unstructured noise. The effect also tended to be stronger (larger effect size) for increasing levels of noise showing that the PW method is less affected by noise in the data. Furthermore, as shown in Figure 3b, the standard method does not efficiently detect oscillator regions with high fixed-ω values (e.g., ω=1.0); on the other hand, it fails to detect a module’s stability with low fixed-ω values (e.g., ω=0.1).

**Figure 3.**
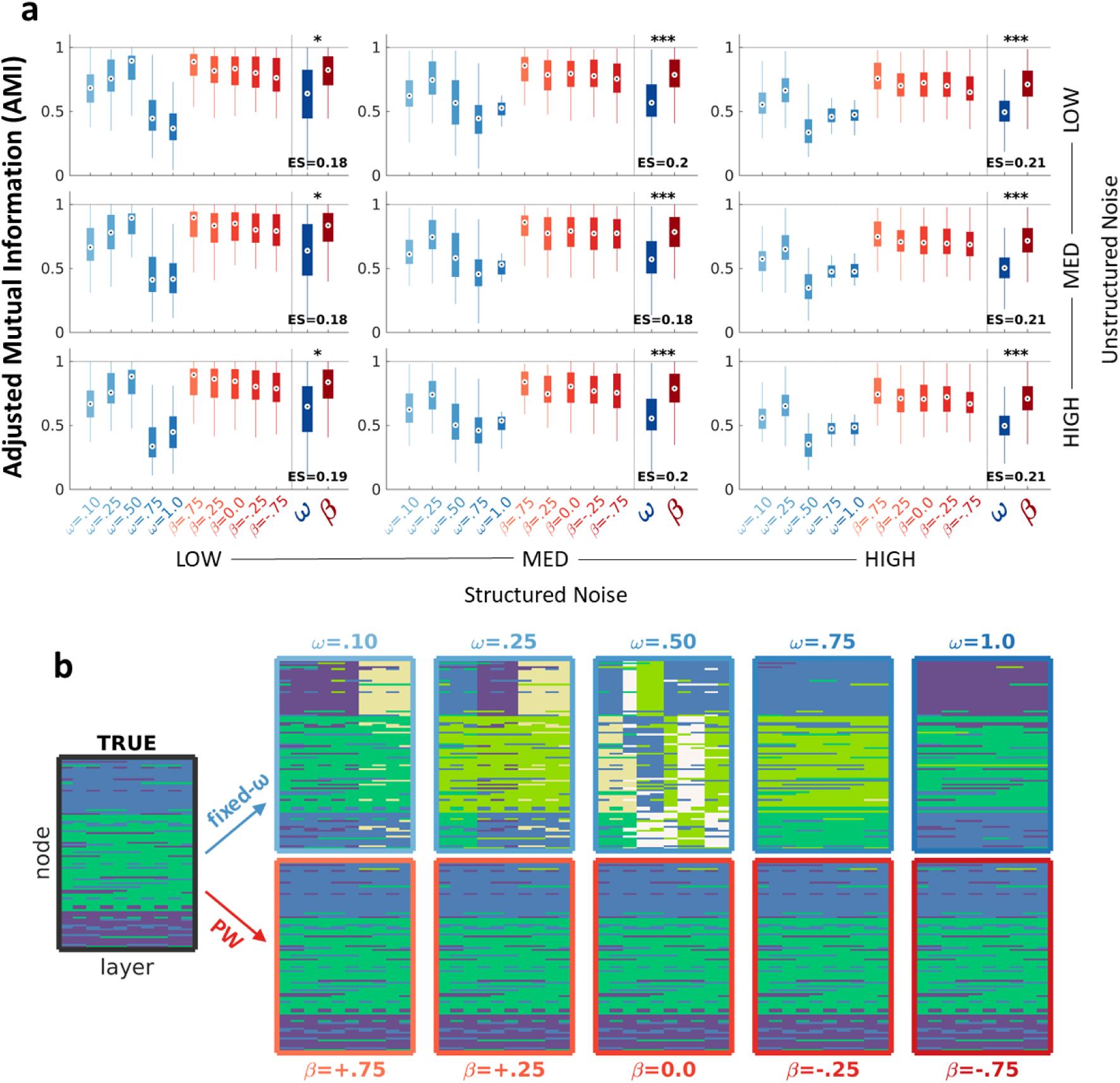
Results from simulated data with a three-modules structure. (a) The PW method (red) outperformed the standard fixed-ω method (blue) as shown by box plot reporting increased adjusted mutual information (AMI). The effect size (ES) for the term *method* in the mixed-effects model indicates, for each combination of noise levels, a performance increase of the PW method in contrast to the fixed-ω method. Significance levels are indicated by the asterisks (***: p<.001, **: p<.01, *: p<.05). (b) Differential community detection for medium noise levels. It is possible to appreciate that the standard method does not efficiently detect oscillator regions with high fixed-ω values (e.g., ω=1.0); on the other hand, it fails to detect module’s stability with low fixed-ω values (e.g., ω=0.1). To note, results reported here are for γ=1.0. For a comprehensive list of cross-γ effects, see Table I.

The same results were obtained with the five-modules structure. Also in this case, the performance of the PW method was better than the performance of the standard fixed-ω method (Figure 4a). Moreover, also in this case the PW method was more resistant to increasing levels of noise. Similar to results for the three-modules structure, the standard method did not efficiently detect oscillator regions with high fixed-ω values and failed to detect module’s stability with low fixed-ω values (Figure 4b).

**Figure 4.**
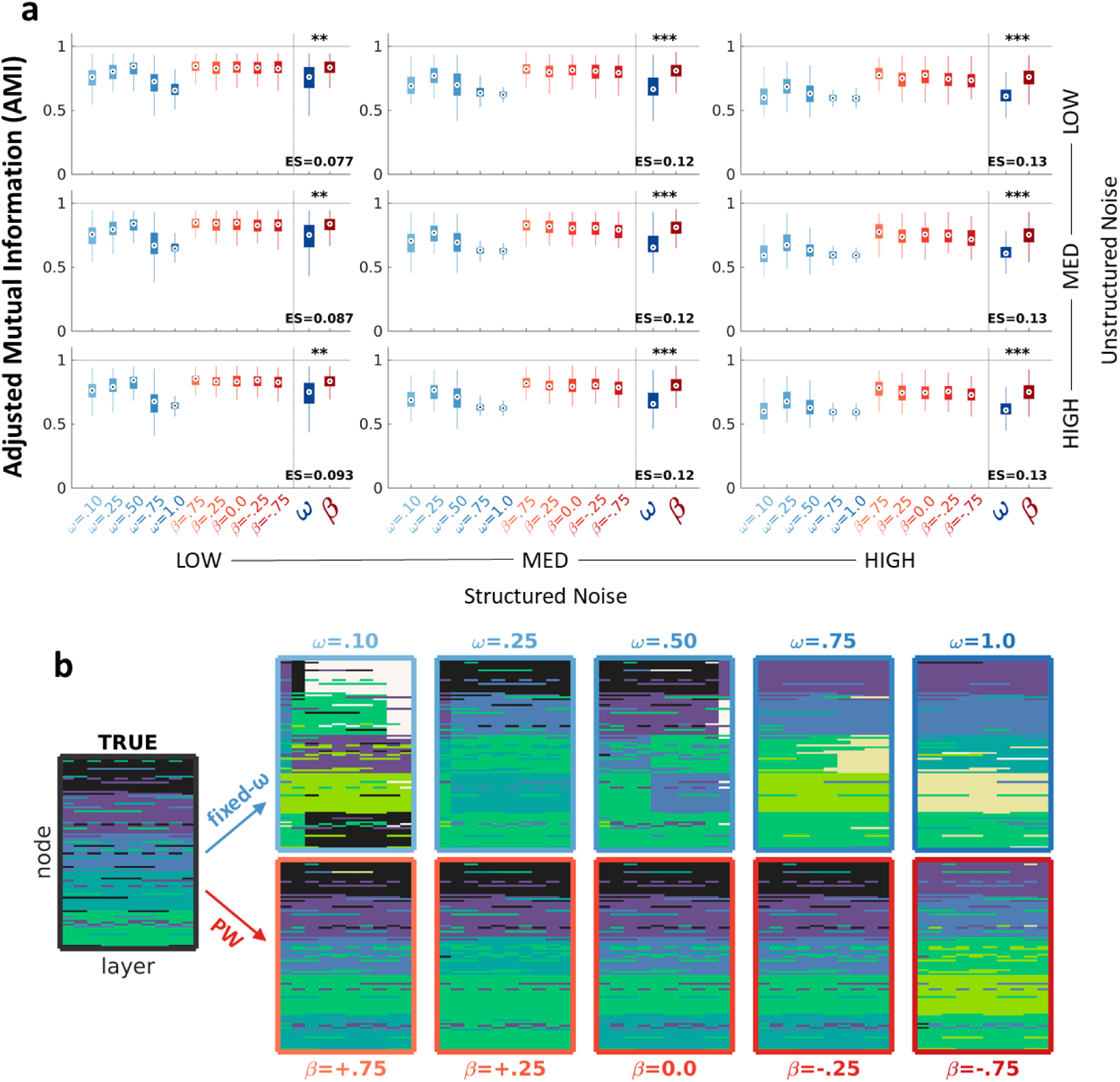
Results from simulated data with a five-modules structure. (a) The PW method (red) outperformed the standard fixed-ω method (blue) as shown by box plots reporting increased adjusted mutual information (AMI). The effect size (ES) for the term *method* in the mixed-effects model indicates, for each combination of noise levels, a performance increase of the PW method in contrast to the fixed-ω method. Significance levels are indicated by the asterisks (***: p<.001, **: p<.01, *: p<.05). (b) Differential community detection for medium noise levels. As in figure 3b, it is possible to appreciate that the standard method does not efficiently detect oscillator regions with high fixed-ω values (e.g., ω=1.0); on the other hand, it fails to detect module’s stability with low fixed-ω values (e.g., ω=0.1). To note, results reported here are for γ=1.0. For a comprehensive list of cross-γ effects, see Table I.

A comprehensive list of cross-γ results is provided in Table I, which reports significance levels and effect sizes (estimates) for the factor *method* across the various simulations in the study. Results for the Rand coefficient are shown in supplementary Figure S1 and in Table S1.

**Table I.** Effects for the fixed term *method* in the linear mixed-effects models investigating the global performance of the PW method versus standard fixed-ω values as indexed by the adjusted mutual information index (AMI). The structural resolution parameter was tuned in the range [0.6 2.0]. Since the table reports the effect size (coefficient) for the term *method* (contrast: PW – fixed-ω), it can be interpreted as the increase in the AMI index when using PW methods instead of fixed-ω methods. It is important to note that the greatest increase in performance coincides with the structural resolution in which the two methods are globally more efficient, showing a considerably better performance in the PW method. It can also be noticed how the effect is greater with higher levels of noise, reflecting the resistance of the PW method (but not of the fixed-ω method) against high noise in the data. Significant effects are highlighted in bold (*: p<.05, **: p<.01, ***: p<.001).

|  | *Unstructured Noise* | **3-modules network** |  |  | **5-modules network** |  |  |
| --- | --- | --- | --- | --- | --- | --- | --- |
|  |  | *Structured Noise* |  |  | *Structured Noise* |  |  |
|  |  | *low* | *med* | *high* | *low* | *med* | *High* |
| $\gamma = 0.6$ | *low* | .10 | **.19\*** | **.20\*\*** | .03 | **.09\*** | **.11\*\*** |
|  | *med* | .11 | **.20\*** | **.20\*\*** | .04 | **.09\*** | **.11\*\*** |
|  | *high* | .12 | **.20\*** | **.20\*\*** | .06 | **.10\*** | **.11\*\*** |
| $\gamma = 0.8$ | *low* | .14 | **.18\*** | **.24\*\*\*** | .05 | .10 | **.12\*\*** |
|  | *med* | .14 | **.18\*** | **.23\*\*** | .06 | .10 | **.12\*\*** |
|  | *high* | .15 | **.18\*** | **.24\*\*\*** | .06 | **.10\*** | **.13\*\*** |
| $\gamma = 1.0$ | *low* | **.18\*** | **.20\*\*\*** | **.21\*\*\*** | .06 | **.09\*** | **.13\*\*\*** |
|  | *med* | **.18\*** | **.18\*\*\*** | **.21\*\*\*** | .06 | **.09\*** | **.14\*\*\*** |
|  | *high* | **.19\*** | **.20\*\*\*** | **.21\*\*\*** | .06 | **.09\*** | **.14\*\*\*** |
| $\gamma = 1.2$ | *low* | **.15\*\*** | **.17\*\*\*** | **.15\*\*\*** | **.07\*** | **.11\*\*\*** | **.13\*\*\*** |
|  | *med* | **.16\*\*** | **.18\*\*\*** | **.16\*\*\*** | **.07\*** | **.11\*\*** | **.13\*\*\*** |
|  | *high* | **.15\*\*** | **.17\*\*\*** | **.16\*\*\*** | **.08\*** | **.12\*\*\*** | **.13\*\*\*** |
| $\gamma = 1.4$ | *low* | **.14\*\*\*** | **.12\*\*\*** | **.09\*\*\*** | **.08\*\*** | **.12\*\*\*** | **.13\*\*\*** |
|  | *med* | **.14\*\*\*** | **.12\*\*\*** | **.09\*\*\*** | **.09\*\*** | **.12\*\*\*** | **.13\*\*\*** |
|  | *high* | **.13\*\*\*** | **.11\*\*\*** | **.09\*\*\*** | **.09\*\*** | **.12\*\*\*** | **.13\*\*\*** |
| $\gamma = 1.6$ | *low* | **.08\*\*\*** | **.05\*\*\*** | **.04\*\*\*** | **.09\*\*\*** | **.11\*\*\*** | **.12\*\*\*** |
|  | *med* | **.08\*\*\*** | **.05\*\*\*** | **.04\*\*\*** | **.10\*\*\*** | **.12\*\*\*** | **.12\*\*\*** |
|  | *high* | **.08\*\*\*** | **.05\*\*\*** | **.04\*\*\*** | **.10\*\*\*** | **.12\*\*\*** | **.12\*\*\*** |
| $\gamma = 1.8$ | *low* | **.04\*\*\*** | **.03\*\*\*** | **.03\*\*\*** | **.09\*\*\*** | **.10\*\*\*** | **.09\*\*\*** |
|  | *med* | **.04\*\*\*** | **.03\*\*\*** | **.03\*\*\*** | **.10\*\*\*** | **.10\*\*\*** | **.09\*\*\*** |
|  | *high* | **.03\*\*\*** | **.03\*\*\*** | **.02\*\*\*** | **.10\*\*\*** | **.10\*\*\*** | **.08\*\*\*** |
| $\gamma = 2.0$ | *low* | **.02\*\*\*** | **.02\*\*\*** | **.02\*\*\*** | **.09\*\*\*** | **.07\*\*\*** | **.06\*\*\*** |
|  | *med* | **.02\*\*\*** | **.02\*\*\*** | **.02\*\*\*** | **.08\*\*\*** | **.07\*\*\*** | **.07\*\*\*** |
|  | *high* | **.02\*\*\*** | **.02\*\*\*** | **.02\*\*\*** | **.08\*\*\*** | **.07\*\*\*** | **.06\*\*\*** |

As expected, results from our simulations indicated that the PW multilayer networks outperformed by far the standard fixed-ω method in detecting true synthetic multilayer networks. The performance achieved with PW multilayer networks on synthetic data was better across structural resolutions and with every teste combination of structured and unstructured noise. Moreover, this better performance using the PW method was permanent using both the three-module and the five-module multilayer structures.

### Real Data

Two significant associations were detected through the analysis of multilayer measures and latent behavioral dimensions. The first one was a positive association between nodal flexibility and factor 2 (F2, psychosis-like experiences). This association was detected by using the standard fixed-ω approach as well as by using PW multilayer modelling. However, the results were not significant for the fixed ω approach with ω=1 (Figure 5a). The analysis of random effects and of individual conditional expectation (ICE) plots showed a consistent heterogeneity of results across nodes (Figure 5b). Nodes with the most prominent effects were located in medial prefrontal, posterior cingulate, and anterior temporal cortices usually ascribed to the default mode network (Figure 5c).

**Figure 5.**
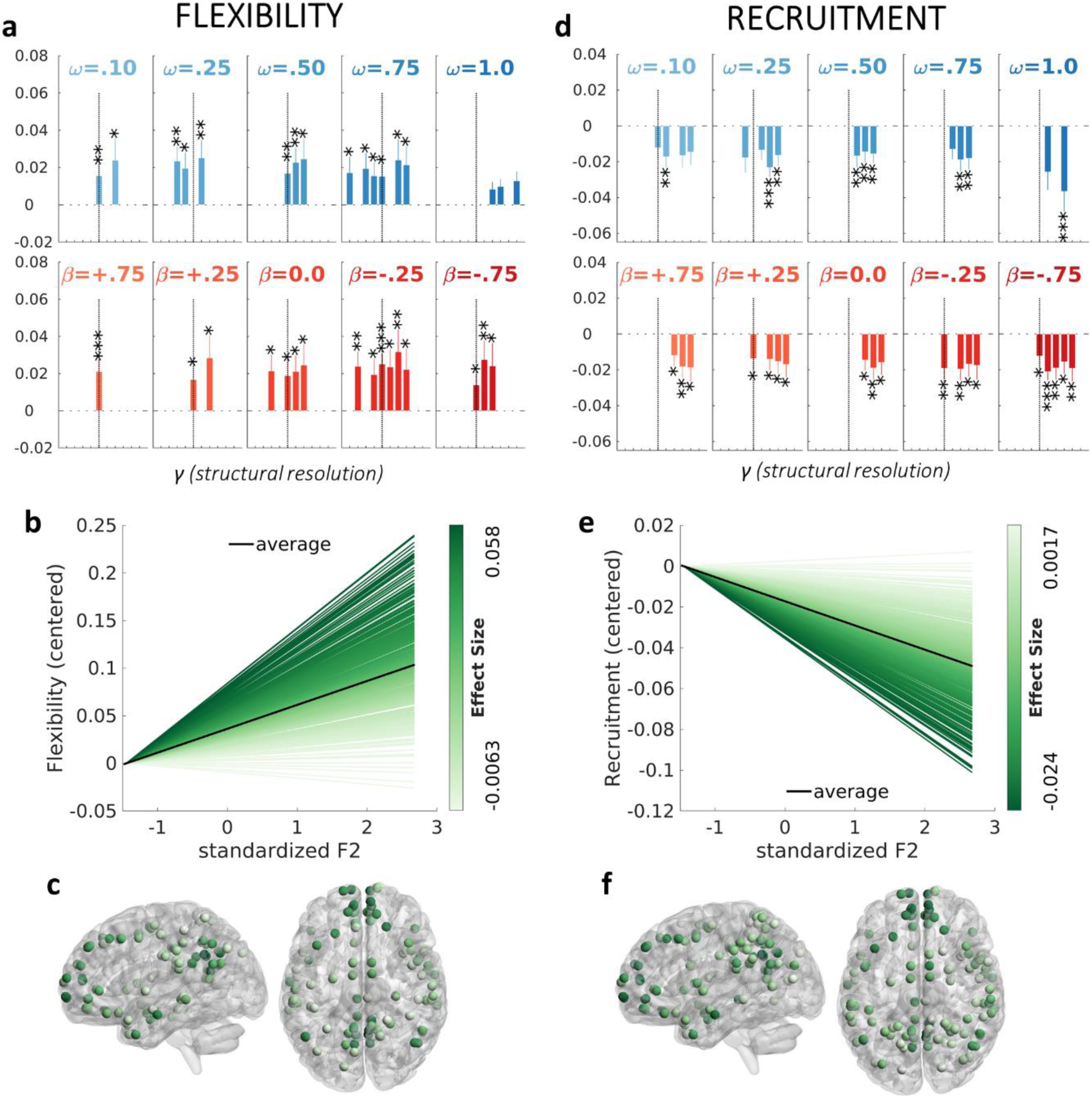
Significant associations between the behavioral factor F2 (psychosis-like experiences) and flexibility (left column) or recruitment (right column). (a) Flexibility was associated with F2 across all the investigated values of β (lower row, red). The association was significant also with four out of five of the fixed-ω values (higher row, blue) but not with ω=1.0. Significance is indicated by the asterisks (***: p<.001, **: p<.01, *: p<.05), whereas the bars without asterisks represent results not significant after correction for multiple comparisons. (b) Individual conditional expectation (ICE) plot representing predictions for flexibility in each brain node with varying levels of psychosis-like experiences (F2). (c) Brain topography of the association between flexibility and F2. Only nodes for which the slope was significantly higher than the average are represented. Colour intensity indicates the strength of the effect size. The subfigures (d-e-f) illustrate the corresponding results for the metric recruitment. For simplicity, only results according to γ=1.0 and to the PW multilayer networks obtained with β=.25 are shown in subfigures b, c, e, and f.

The second significant effect concerned a negative association between nodal recruitment and factor 2. The effect was detected with both the standard fixed-ω method and the PW multilayer networks (Figure 5d). Also in this case, there was a high degree of heterogeneity of effect sizes across brain regions (Figure 5e) and nodes primarily involved were located in anterior prefrontal, posterior cingulate, and anterior temporal regions of the default mode network (Figure 5f).

No significant results were observed regarding promiscuity or integrity.

## DISCUSSION

The primary aim of this study was to test the performance of the probabilistically weighted (PW) method for detecting cross-layer communities. We implemented this method both in synthetic, perturbated time series of a network and in real data from resting-state fMRI. When applied to synthetic data, the PW method outperformed the standard method and was less affected by increasing noise levels in the time series. The better performance of the PW method was confirmed using the adjusted mutual information index and the Rand coefficient. Moreover, a better performance was found for the PW method using modular structures with an increasing number of modules. Importantly, standard fixed-ω approaches did not efficiently detect oscillator regions with high fixed-ω values (e.g., ω=1.0) and failed to detect modular stability with low fixed-ω values (e.g., ω=0.1).

The improvement in accuracy brought by our PW method was consistent across different structural resolution (γ) values. These results are the direct consequence of a reduced bias of selection with respect to the strength of cross-layer intra-nodal connections in the PW method. When applied to real data, the PW method was able to identify significant associations of two dynamic metrics, namely flexibility and recruitment, with behavioral measures. We postulate that probabilistically weighted multilayer networks are desirable for implementation in future studies and should replace the standard, biased fixed-ω method.

To note, the method presented in this paper is intrinsically developed, and thus more suitable, for weighted networks. However, it can also be applied to binarized graphs since the intra-nodal weights can be directly transformed in a stochastic binary connection value. Of course, binarizing cross-layer connections makes sense in a framework in which also intra-layer functional connectivity is binarized. However, the focus of the present study is not on binarized versus weighted procedures for network analysis.

The results on real resting-state fMRI data also deserve further discussion. Flexibility indicates how frequently a node changes module allegiance over time. Instead, recruitment indicates the probability of a node to belong to the same community as other nodes from its own system. Thus, our results indicate that a decreased stability of nodes within the DMN correspond to higher levels of psychosis-like experiences (F2) in healthy adults. These findings are in line with the role of the fragmentation of the DMN in the predisposition toward psychopathology involving psychosis and dysfunctional behavior (Braun et al., 2016; Hu et al., 2017; Hua et al., 2019; Fan et al., 2019). Moreover, higher flexibility and less stable modular organization have already been shown in patients with schizophrenia (Braun et al., 2015; Gifford et al., 2020), whereas higher switching has been associated with impaired sleep and decreased behavioral performances (Thompson et al., 2018; Pedersen et al., 2018). In this framework, our findings also fit with DMN-specific network aberrancies observed in patients with psychotic disorders (Ebisch and Aleman, 2016; Du et al., 2018; Di Plinio et al., 2020a; Humpston and Broome, 2020), which also have genetic underpinnings (Scariati et al., 2014). We contribute to this framework by showing for the first time an association between nodal instability within the DMN (especially in medial prefrontal and in posterior cingulate cortices) and psychosis-like experiences in a sample of healthy individuals. We remark that these findings were possible due to the implementation of the PW method here presented. Considering that an aberrant degree of psychosis-like experiences may imply difficulties in social interactions (De Bézenac et al., 2015) and eventually impaired social functioning (Nelson et al., 2012), future studies are needed to assess how the decreased stability of self-networks (DMN) may predispose toward clinical self-disturbances in healthy populations (Nelson et al., 2012; Orr et al., 2014; Humpston et al., 2014; McGrath et al., 2015).

When constructing and analyzing brain activity and functional connectivity, ground truths are rare and unstable. However, advances in the understanding of dynamic brain networks point toward the existence of multi-scale entities in the brain architecture (Hutchinson et al., 2013; Boccaletti et al., 2014; Kivela et al., 2014; Braun et al., 2015; Telesford et al., 2016; De Domenico et al., 2017; Betzel and Bassett, 2017; Zheng et al., 2018; Thompson et al., 2018; Betzel et al., 2019; Gifford et al., 2020; Yang et al., 2021; Tardiff et al., 2021). These advances followed recent developments in network science (Newman, 2003; Rubinov and Sporns, 2010; Bassett and Sporns, 2017), and generally indicate the necessity to implement unbiased methods for detecting communities (Newman and Girvan, 2004; Porter et al., 2009; Fortunato, 2010; Lancichinetti et al., 2011; Fenn et al., 2012; Sporns and Betzel, 2016; Fortunato and Hric, 2016), possibly controlling for multiple stochasticity in the community organization, like in multi-resolution approaches (Rubinov et al., 2015; Chai et al., 2016; Puxeddu et al., 2019; Di Plinio et al., 2020a; Malagruski et al., 2020). In fact, the choice of the resolution parameter γ is important since it regulates the number and size of communities in a network (Lancichinetti and Fortunato, 2009, 2012; Betzel and Bassett, 2017). However, in multilayer networks, the weight of intra-nodal (cross-layer) links become crucial if we are interested in temporal features of the network such as flexibility, recruitment, or promiscuity (Bassett et al., 2011; Papadopouolos et al., 2016; Telesford et al., 2017; Pedersen et al., 2018). The implementation of PW multilayer networks allows an unbiased approach to the study of brain networks through multiple temporal resolutions and without a-priori guessing the strength of cross-layer connections.

Concluding, in the present study we introduce a probabilistic, biologically driven approach for the unbiased selection of cross-layer weights in multilayer networks. Crucially, our study does not only propose a way of optimizing cross-layer functional connections for accurate modular detection, but also renovate the theoretical conception of multilayer brain networks, favouring the shift toward a probabilistic and biologically-sound translation of timelines to modules. Probabilistic multilayer networks allow the proper study of multiple temporal resolutions, represented by multiple shape (β) values in the distribution of intra-node connection weights. The PW method performs optimally despite modulations of β (probabilistic weights), indicating that it can model a plurality of multilayer networks without suffering from the biases previously described from fixed-ω approaches. Furthermore, it investigates multiple temporal resolutions in multilayer networks that are biologically grounded since they are assigned based on the coherence across time windows, paralleling the actual unbiased approach commonly employed for spatial resolution (γ). However, we suggest to rely on the study of several values of probabilistic connections including many β values in the analysis (or equivalent modelling of probabilistic cross-layer weights). The ground truth in the brain functioning is complex and cannot be achieved by searching for a single optimized value since the brain probably functions at many levels of structural resolutions simultaneously. Thus, we suggest relying on the analysis of multiple γ and β parameters to favour unbiased research. In fact, as we shown in the present study, the use β modulations indeed combines well with the use of multiple γ. The higher performance achieved by the PW method, together with the significant associations with psychotic-like experiences detected, is a step forward towards an unbiased approach in the study of dynamic brain functioning as well as its behavioral and cognitive correlates.

